# Stat3 has a different role in axon growth during development than it does in axon regeneration after injury

**DOI:** 10.1101/2022.12.05.519245

**Authors:** Qinwen Duan, Hongfei Zheng, Yanjun Qin, Jizhou Yan, Shawn Burgess, Jian Wang, Chunxin Fan

## Abstract

Signal transducer and activator of transcription 3 (STAT3) is essential for neural development and regeneration as a key transcription factor and mitochondrial activator. In this study, using zebrafish posterior lateral line (PLL) axons, we demonstrate that Stat3 plays distinct roles in PLL axon embryonic growth and regeneration. In zebrafish *stat3* mutants the PLL axons were truncated during embryonic growth. Expressing *stat3* in neurons of the PLL ganglion rescued their axon growth suggesting *stat3* promoted PLL axon growth in a cell-autonomous manner. Using Jak/Stat signaling inhibitors or modified Stat3, we found Jak/Stat signaling was dispensable for PLL axon growth. We showed that Stat3 was co-localized with mitochondria in PLL axon and lower ATPase activity in *stat3* mutant embryos than wild-type embryos. The PLL axon growth was affected by mitochondrial modulators indicating that Stat3 regulated PLL axon growth through mitochondrial Stat3 function. Mutation of *stat3* or treating with Jak/Stat signaling inhibitors retarded PLL axon regeneration and Schwann cell migration. Taken together, Stat3 is required for embryonic PLL axon growth by regulating the ATP synthesis efficiency of mitochondria, whereas Stat3 stimulates PLL axon regeneration through regulating Schwann cell migration via Jak/Stat signaling. Our findings show a new mechanism of Stat3 in axon growth and regeneration.

**Author Summary:** Neurons are one of the most specialized cell type in our body. A slender axon used for transmitting information between neuron and target cell extends from the cell body. Axon extension during both development and regeneration is regulated by intrinsic and extrinsic factors. In this study, we used the zebrafish posterior lateral line (PLL) axons which extends a long distance to study the regulation of axon extension. Signal transducer and activator of transcription 3 (STAT3) is a key transcription factor in Jak/Stat signaling and an activator for mitochondria functions. Our results demonstrate that Stat3 plays distinct roles in PLL axons development and regeneration. During embryonic development, Jak/Stat signaling is dispensable for PLL axons extension, whereas the mitochondrial Stat3 is required for embryonic axon growth through promoting ATP synthesis. During PLL axons regeneration, Jak/Stat is essential for Schwann cells migration which mediates axon regeneration.

## Introduction

Axon extension is a critical process for both embryonic neuronal development and regeneration after injury. The extension of axons requires both intrinsic and extrinsic factors (Kaplan et al., 2015). During axon extension, significant energy is required for forming the abundant dynamic actin filaments in growth cones, microtubule-based axonal transport of organelles and proteins, and synthesis of protein and phospholipids used for membrane expansion. Oxidative phosphorylation (OXPHOS) in mitochondria is the main source of ATP synthesis. Recent investigations have shown a role for mitochondria in axon extension (Sheng, 2017; Smith and Gallo, 2018). Besides providing energy, axon regeneration requires additional processes such as axon debris clearance, scar gap bridging, and guiding the migration of axons. All of these processes require glia cells wrapping around the regenerating axons (Min et al., 2021).

Signal transducer and activator of transcription 3 (STAT3) is a conserved protein with multiple essential functions (Avalle and Poli, 2018). The most widely studied function of Stat3 is in mediating the signaling from extracellular cytokines and growth factors. Stat3 is phosphorylated by Janus kinase (JAK) and other tyrosine kinases at a tyrosine in the SH2 domain, after which it dimerizes and translocates to the nucleus to activate genes involved in inflammation, cell proliferation, and/or survival (Kiu and Nicholson, 2012). Recently, Stat3 was also found to bind to complex I and II of ETC, and ATP synthase in the inner mitochondrial membrane and to promote the electron transport chain (ETC) and ATP synthesis (Gough et al., 2009; Rincon and Pereira, 2018; Wegrzyn et al., 2009; Yu et al., 2014).

Some studies provide evidence that Stat3 is involved in axon extension through Jak/Stat signaling. Deletion of suppressor of cytokine signaling 3 (SOCS3), an inhibitor for Jak, promotes robust regeneration of injured optic nerve axons in mice (Smith et al., 2009). In contrast, over-expression of *socs3* attenuates optic nerve regeneration in zebrafish (Elsaeidi et al., 2014). Nerve regeneration is also regulated by the activation of the Jak/Stat pathway in Schwann cells (Benito et al., 2017; Lin et al., 2016). However, other studies have shown that mitochondrial Stat3 is essential for axon extension. Mitochondrial Stat3 promotes neurite outgrowth in PC12 cells and cultured dorsal root ganglion (DRG) neurons (Hwang and Namgung, 2021; Zhou and Too, 2011). In mouse retinal ganglion cells (RGC), fusing Stat3 with a mitochondria targeting sequence (MTS) promotes further enhancement in axon regeneration compared to the constitutively active form of Stat3 (Luo et al., 2016).

The lateral line is a specific sensory system present in fish and amphibian. The posterior lateral line (PLL) ganglion is located posterior to the otic vesicle and relays information from the lateral line sensory organs, known as neuromasts, to the medial octavolateralis nucleus (MON) in the hindbrain. During embryonic development, the axons of the PLL nerves extend from the ganglion to the tip of tail over the course of approximately 20 h (Pujol-Martí and López-Schier, 2013). In addition, the PLL axons can regenerate spontaneously after neuroaxotomy (Villegas et al., 2012). This makes the PLL an ideal model system for axonal growth studies in vivo (Ceci et al., 2014). Stat3 is expressed specifically in lateral line, including the neuromasts and lateral line ganglia (Liang et al., 2012; Liu et al., 2017). In this study, we generated a zebrafish *stat3* mutant and showed its PLL axons failed to extend properly. We identified a cell-autonomous role of Stat3 in PLL axon growth through a PLL neuron specific rescue assay. Unexpectedly, we found that mitochondrial Stat3 rather than Jak/Stat signaling was required for PLL embryonic axonal growth. In contrast, Jak/Stat signaling inhibitors retarded PLL axon regeneration. Thus, Stat3 plays distinct roles in PLL axon embryonic growth and regeneration.

## Results

### 1. stat3 is required for PLL axon growth in the zebrafish embryo

It has been shown in previous studies that *stat3* is enriched in the cranial ganglion and lateral line system during the larval stages (Liang et al., 2012; Liu et al., 2017). We examined the expression of *stat3* during lateral line axon growth and found that *stat3* was enriched in anterior lateral line (ALL) and PLL at 32 hpf, disappearing after 48 hpf (Fig 1A). The dynamic expression of *stat3* in lateral line ganglion suggests that it is probably involved with the early formation of the lateral line neurons.

**Fig 1.**
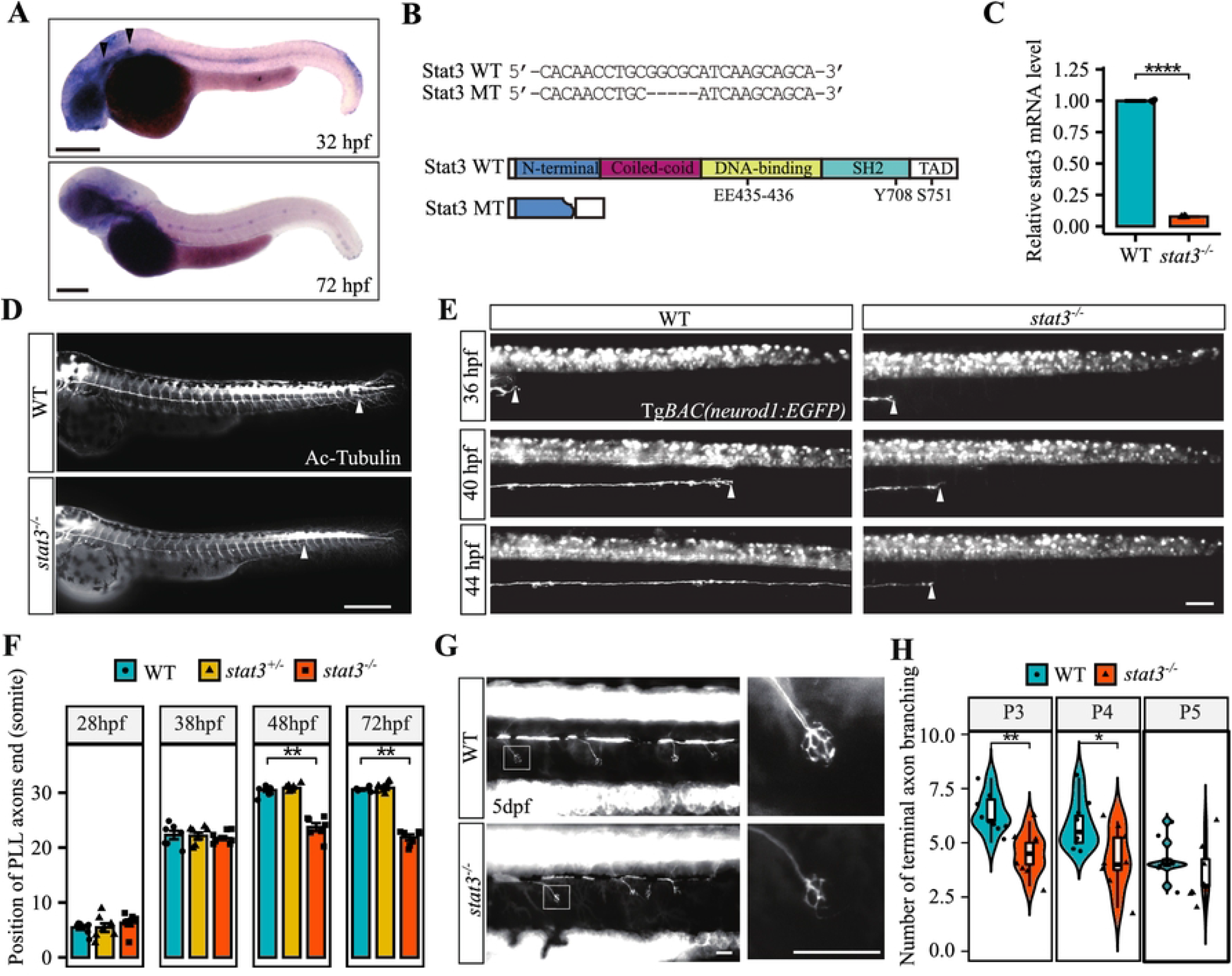
*stat3* is essential for posterior lateral line (PLL) axon growth. (A) *In situ* hybridization shows *stat3* is mainly expressed in lateral line ganglion and neuromasts at 32 hpf and 48 hpf. Arrows mark the ALL and PLL ganglion. Scale bars are 200 μm (B) The effect of zebrafish *stat3* mutant on Stat3 DNA and protein sequence. Alignment of the *stat3* homozygous mutant allele with the corresponding wild-type sequence showing the deleted nucleotides (top panel). The functional domains in wild-type Stat3 protein (806 AA) and the putative truncated protein (167 AA) (bottom panel). (C) Quantification of *stat3* mRNA expression in *stat3* homozygous mutants relative to wild-type at 2 dpf using qPCR (3 replicates). **** *p* < 0.0001 (*t*-test). Error bar indicates SEM. (D) PLL axons were immune-stained with anti-acetylated-tubulin antibody in wild-type and *stat3* mutants at 2 dpf. Arrows point to the posterior end of PLL axons. Scale bar, 500 μm. (E) Lateral views of the representative PLL axon outgrowth process in wild-type and *stat3* mutant are shown using TgBAC*(neurod1:EGFP)*. The arrows point to the posterior end of PLL axons. Scale bar, 100 μm. (F) Quantification of the PLL axon length shown as the somite level for the posterior end of PLL axons *(N* = 7). ** adjusted *p* < 0.01 (Wilcoxon RS test), compared with wild-type. Error bars indicate SEM. (G) PLL axon terminals that innervate neuromasts were visualized using TgBAC*(neurod1:EGFP)* in wild-type and *stat3* mutants at 5 dpf (left panel). The right panels are magnifications of the terminal branches. Scale bar, 50 μm. (H) Quantification of the number of terminal branches in PLL axons of *stat3* mutants and wild-types (*N* = 8). * *p* < 0.05 (Wilcoxon RS test), compared with wild-type. Error bar indicates SEM.

To study the role of *stat3* in PLL ganglion development, we knocked-out *stat3* in zebrafish using CRISPR/Cas9 and created a zebrafish *stat3* mutant line with 5 bp-deletion resulting in a premature stop codon. The putative truncated protein has an incomplete N-terminal domain and is missing the coiled-coil, DNA-binding, SH2 (Src Homology 2), and transcription activation domains (TAD) compared to wild-type Stat3 (Fig 1B). qPCR results indicated that wild-type the expression of *stat3* is significantly decreased in the *stat3* homozygous mutants, which suggested nonsense-mediated mRNA decay was induced by the mutation (Fig 1C). We also found *stat3* mutants had curved bodies at approximately 20 dpf (data not shown) as described previously (Liu et al., 2017). These results indicated that *stat3* was successfully knocked-out in the mutant line.

To determine the effect of a *stat3* mutation on the PLL ganglion, we labeled axons using anti-acetylated-tubulin antibodies. By 48 hpf, the PLL axons reached the caudal neuromasts in wild-type. However, the PLL axons in *stat3* mutants only migrated approximately 70% of the distance of wild-type siblings (Fig 1D). We crossed TgBAC*(neurod1:EGFP)* transgenic zebrafish, which labels both the cell body and axon of lateral line ganglion with enhanced green fluorescent protein (EGFP) (Obholzer et al., 2008), with *stat3* mutation to enable tracing PLL axon growth using time-lapse imaging. The initiation of PLL axons occurred at approximately 20 hpf for both the *stat3* mutant and wild-type siblings. Growth speed didn’t show significant differences by 20-38 hpf, however, the pioneer growth cone of PLL axons stopped migrating and got thinner in *stat3* mutants at about the 20^th^-22^nd^ somite while the PLL axons reached to caudal neuromasts in wild-type (N=7) (Fig 1E-F; Video S1A-B). We also checked the PLL axons in *stat3* mutants after 2 dpf. Although the PLL axons in *stat3* mutants became thinner during migration, they didn’t show obvious degeneration or retraction even by 17 dpf (S1 Fig.).

To elucidate whether axon truncation in *stat3* mutants was due to loss of neurons innervating the caudal neuromasts specifically, we counted the cell bodies in PLL ganglion in the TgBAC*(neurod1:EGFP)* transgenics at 30 hpf. The PLL ganglion in *stat3* mutant contained similar average number of neurons (22.0 ± 0.97) compared to wild-type (22.3±0.63) (S2 Fig.). The *stat3* mutation does not affect the number of cell bodies in the PLL ganglion.

In addition, we counted the terminal branches under TgBAC*(neurod1:EGFP)* transgenic background and the *stat3* mutants displayed fewer terminal branches in PLL axons innervating the trunk neuromasts, however, it didn’t show the swollen axon terminals seen in the *jip3^nl7^* mutant described previously (Fig 1G-H) (Drerup and Nechiporuk, 2013). These data suggest that the underlying mechanism of PLL axon truncation in *stat3* mutants is mechanistically different from the *jip3^nl7^* mutants.

The axons of the dorsal lateral line (DLL) also project from the PLL ganglion. We found the length of DLL axons in *stat3* mutant was also significant shorter than wild-type, and the branch of DLL in *stat3* mutant was fewer than wild-type (S3A Fig.). Blocking Jak/Stat signaling with morpholinos targeting *stat3* and *jak1* affected pathfinding and axon branching in spinal nerves (Conway, 2006). However, in *stat3* mutants, neither the ventral nor the dorsal projections of the spinal nerve were significantly changed (S3B Fig.). There is no obvious expression of *stat3* in spinal nerves so this would be an expected result.

### 2. Stat3 promotes PLL axon growth in a cell-autonomous manner

To illustrate whether Stat3 is involved in Schwann cells differentiation during embryonic development, we analyzed the expression of *sox10*, a marker of neural crest, using *in situ* hybridization in the *stat3* mutants before PLL axons stopped migrating at 30 hpf. In both *stat3* mutants and wild-type, *sox10* was expressed in the anterior region of the horizontal myosyptum (Fig 2A), which indicated that Schwann cell precursors were not affected by the *stat3* mutation. We crossed the *stat3* mutation into the Tg*(mbp:EGFP)* transgenic line whose Schwann cells are labeled with EGFP (Jung et al., 2010) and analyzed mature Schwann cells in the *stat3* mutants. At 3 dpf, the Schwann cells in *stat3* mutants formed normally at the anterior part of the PLL, however they were sparse at the posterior part of PLL (Fig 2B). We analyzed the ultrastructure of PLL neural fibers using transmission electron microscopy between the third and sixth somites at 72 hpf. PLL axons in both the wild-type and *stat3* mutants were correctly myelinated. The thickness of the myelin showed no obvious difference between mutants and wild-types. However, the PLL axons in *stat3* mutants were thinner than wild-type (Fig 2C-D), which confirmed the observations in the TgBAC*(neurod1:EGFP)* transgenic. The decrease in myelination in *stat3* mutant PLL is likely the result of PLL axons failing to extend properly.

**Fig 2.**
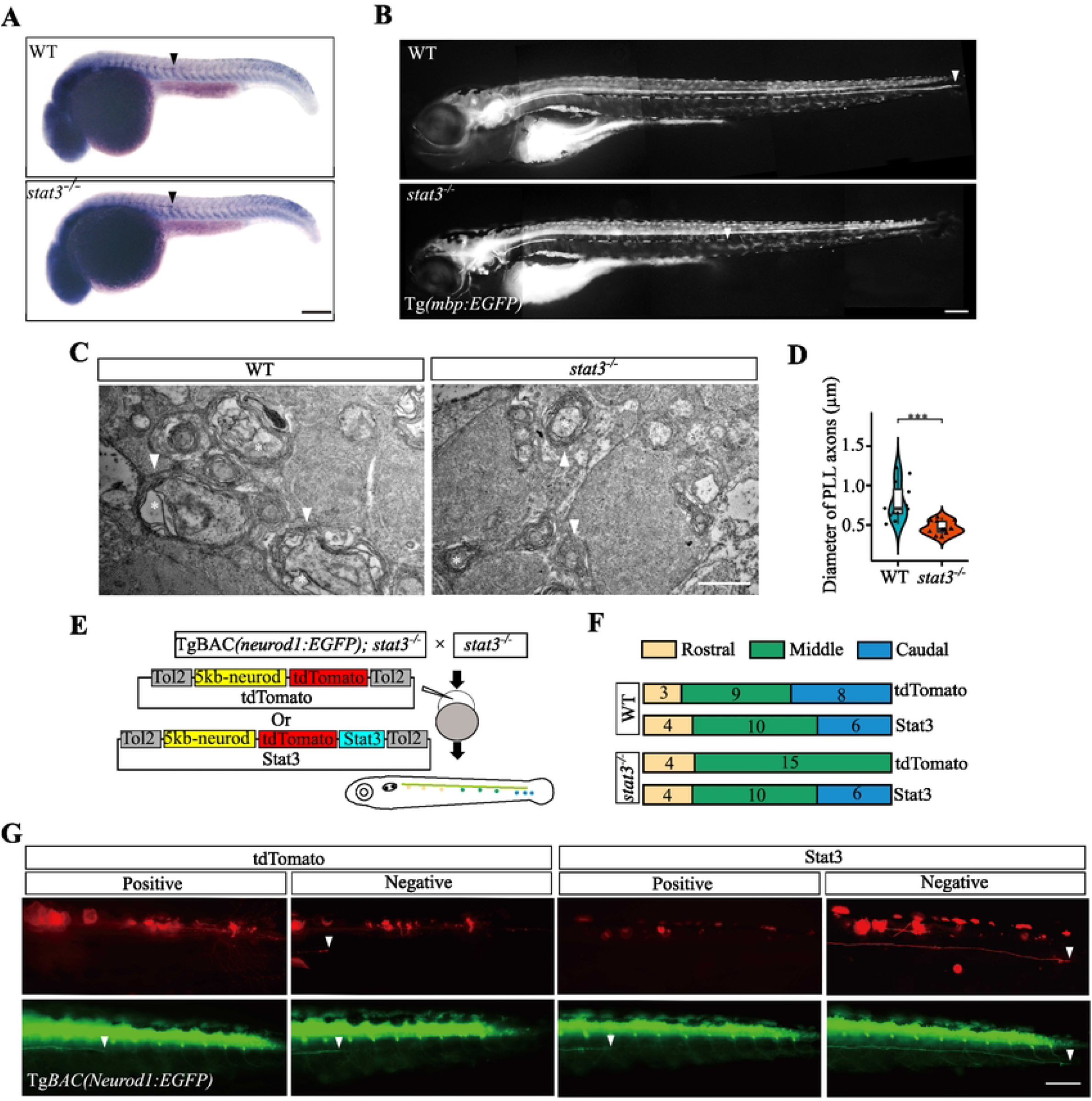
*stat3* promotes PLL axon growth in a cell-autonomous manner. (A) The expression of *sox10* in Schwann cell precursors (arrow) in *stat3* mutant and wild-type at 30 hpf. Scale bars: 100 μm. (B) The Schwann cells around PLL axons visualized by Tg*(mbp:EGFP)* in wild-type and *stat3* mutant at 3 dpf. Scale bar: 100 μm. (C) Transmission electron microscope (TEM) images of transverse sections between P1 and P2 at 3 dpf showing the ultrastructure of the PLL axons in a *stat3* mutant compared to wild-type. Black stars indicate axons, and arrows label the myelin sheath. Scale bar: 0.5 μm. (D) Quantification of PLL axon diameters. *N* = 12. *** *p* < 0.001 (*t*-test). Error bar indicates SEM. (E) Schematic of the rescue assay for expressing of tdTomato-tagged Stat3 or tdTomato (control) in some neurons of the PLL ganglion using transient transgenic constructs. (F) The frequency of PLL tdTomato labeled axons reaching the anterior (P1-P3), middle (P4-P6), and caudal (P7-P9) regions in wild-type and *stat3* mutants after injecting with transient transgenic constructs. (G) Representative results of the rescue assay using Stat3 and Control constructs in *stat3* mutants. The PLL axon truncation phenotype in *stat3* mutants was shown by TgBAC*(neurod1:EGFP)* (green). The PLL axon expressing tdTomato in labeled side was truncated, whereas some PLL axon expressing tdTomato-Stat3 reached to the tip of tail. Arrowheads point to posterior end of PLL axon. Scale bars: 100 μm.

We asked whether *stat3* was involved in PLL axon growth in a cell-autonomous manner or cell-non-autonomous manner. We used the construct encoding tdTomato tagged Stat3 driven by the 5 kb-neurod to express *stat3* in some neurons of PLL ganglion (Neurod-Stat3). The construct only containing the reporter gene, tdTomato was used as a control. Either the Neurod-Stat3 or a tdTomato construct was injected into the offspring of a *stat3*;TgBAC*(neurod1:EGFP)* heterozygous carrier incross. Each larva was imaged to determine the growth of the PLL axons and then genotyped. EGFP or tdTomato fluorescence was used to show the PLL axon phenotype and neurons with *stat3* overexpression, respectively (Fig 2E). The labeled PLL axons were divided into rostral, middle or caudal according to which neuromast they innervated. In wild-types injected with Neurod-Tomato and Neurod-Stat3, similarly labeled PLL axons reached caudal neuromasts. This indicated that extra exogenous Stat3 did not affect PLL axon growth. In *stat3* mutants injected with Neurod-Tomato, no PLL axons reached caudal neuromasts (0/19), however, Neurod-Stat3 extended the PLL axons to caudal significantly (6/20), Fisher’s exact test *p*=0.0316 (Fig 2F). Fig. 2G shows the representative *stat3* mutant larvae injected with the rescue constructs. On the side with or without Neurod-tdTomato labeling, the PLL axons remained truncated. On the side with Neurod-Stat3 labeling, some PLL axons reached the caudal neuromasts (Fig 2G). These results demonstrate that Stat3 is required autonomously for PLL axon extension.

### 3. Jak/Stat signaling is dispensable for PLL axon embryonic growth

The main function of Stat3 is the transcriptional regulation of target genes, which are involved in cell differentiation, migration, and proliferation. Using qPCR, we found that the expression of most Stat3 target genes were decreased significantly in zebrafish *stat3* mutants (S4 Fig.). As reported, Jak/Stat signaling is important for axon regeneration in mouse and zebrafish retinal ganglion cells (RGC) (Elsaeidi et al., 2014; Smith et al., 2009). To investigate whether PLL axon growth is regulated by Jak/Stat signaling, we treated the wild-type zebrafish embryos with Jak/Stat signaling inhibitors during PLL axon development (24 hpf-48 hpf). Stattic inhibits tyrosine phosphorylation of Stat3 (Schust et al., 2006), Pyridone 6 is a pan-JAK inhibitor (Pedranzini et al., 2006), and S3I-201 inhibits Stat3-dependent transcription by blocking Stat3 dimerization and Stat3 DNA-binding activity (Siddiquee et al., 2007). Unexpectedly, the length of PLL axon was not changed significantly after treated with above inhibitors (*N*=20) (Fig 3A-B).

**Fig 3.**
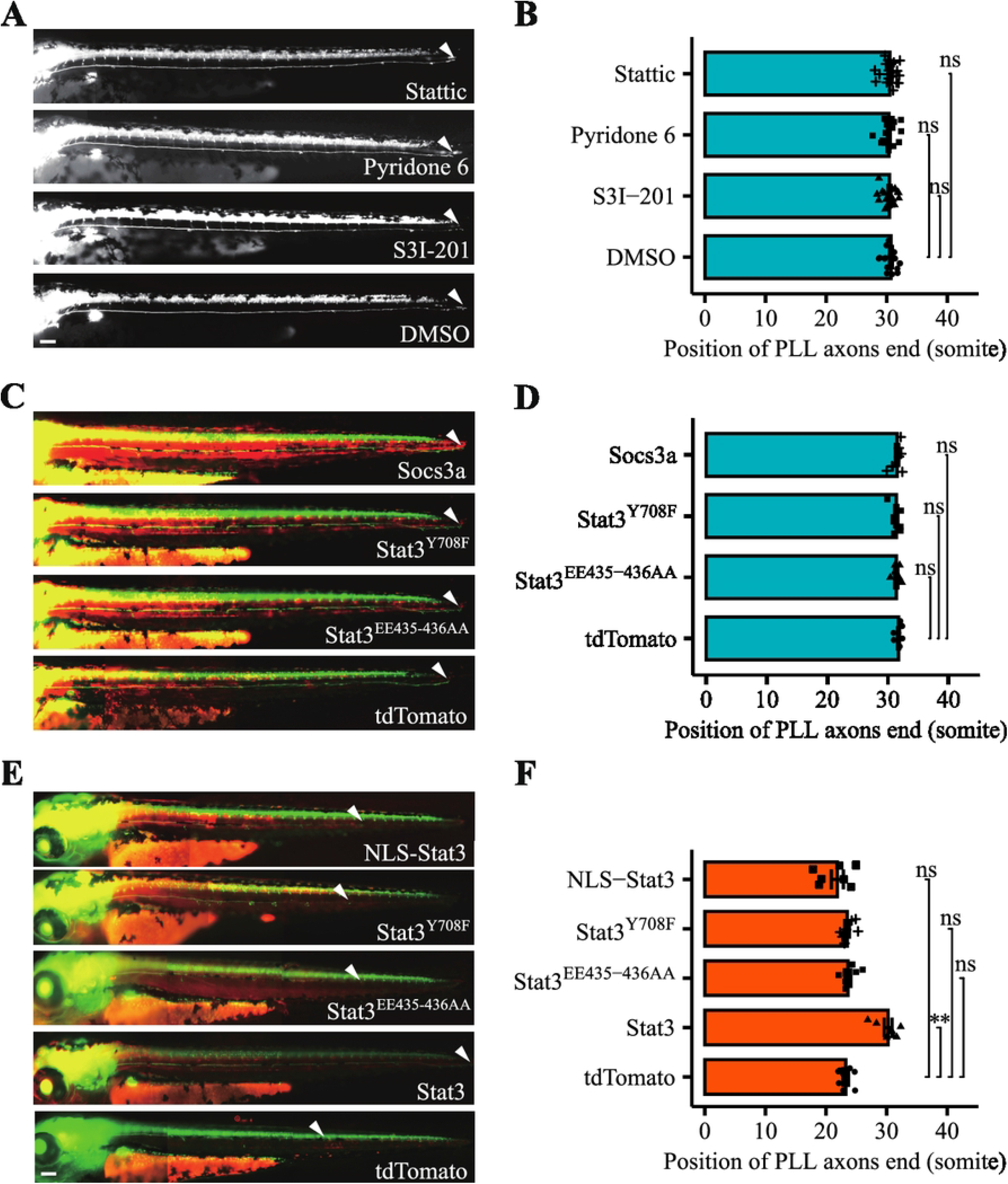
PLL axon growth is independent of Jak/Stat signaling. (A) PLL axons of the TgBAC*(neurod1:EGFP)* transgenic line after treating with Jak/Stat signaling inhibitors (S3I-201, Pyridone 6 or Stattic). DMSO was used as control. (B) Quantification of PLL axon length after treating with Jak/Stat signaling inhibitors. *N* = 20. (C) PLL axons of the TgBAC*(neurod1:EGFP)* transgenic line injected with Jak/Stat signaling blocking mRNA of *socs3a*, Stat3^Y708F^, Stat3^EE435-436AA^. tdTomato mRNA was used as the control. (D) Quantification of PLL axon length after mRNA injections. *N* = 8. (E) PLL axons of *stat3* mutants in the TgBAC*(neurod1:EGFP)* transgenic background rescued with wild-type, mutated, or nuclear localization sequence tagged (NLS) Stat3. (F) Quantification of PLL axon length in *stat3* mutants after mRNA injections. *N* = 8. Arrowheads point to posterior end of PLL axons. Scale bars: 100 μm. The length of PLL axon is quantified by the somite level of posterior end of PLL axons. tdTomato mRNA was used as the control. Somite level differences were examined between each experimental and control group using Wilcoxon Rank Sum test and multiple correction. ** adjusted *p*<0.01 and NS represents no significant difference, compared with control group. Error bar indicates SEM.

A Jak inhibitor and a dominant negative Stat3 were also used as an additional approach to inhibit Jak/Stat signaling and to confirm pharmacological results. We injected zebrafish Socs3a mRNA (an inhibitor of Stat3) into fertilized egg and found the PLL axons could still reach caudal neuromasts at 48 hpf. The substitution of double glutamic acids in the Stat3 DNA binding domain to alanines (EE434-435AA) or tyrosine in the SH2 domain to phenylalanine (Y705F) leads to a dominant-negative effect on Stat3 function (Conway, 2006; Selvaraj et al., 2012). However, both the injection of Stat3^Y708F^ and Stat3^EE435-436AA^ mRNA did not affect PLL axon growth (*N*=8) (Fig 3C-D). All these results suggested Jak/Stat signaling is not essential for zebrafish embryonic PLL axon growth.

We further injected wild-type or modified *stat3* mRNA into *stat3* mutant zebrafish embryos to investigate which function of Stat3 is necessary for PLL axons growth. The growth of PLL axons in *stat3* mutants could be rescued by injecting wild-type *stat3* mRNA. However, a Stat3 containing a strong nuclear localization signal (NLS-Stat3) did not rescue PLL axon growth in *stat3* mutants. The modified Stat3^Y708F^ and Stat3^EE435-436AA^ also failed to rescue axonal growth (N=8) (Fig 3E-F). This indicated that nuclear Stat3 is not sufficient for PLL axon growth, but the SH2 and DNA binding domains are still required for its function in PLL axon growth.

### 4. Mitochondrial Stat3 regulates embryonic PLL axon growth by promoting ATP synthesis

As reported, a small amount of Stat3 localizes to the mitochondria and enhances electron transport chain (ETC) function and increase ATP synthesis in mammalian cell lines (Wegrzyn et al., 2009). We asked whether mitochondrial Stat3 was involved in PLL axon growth in zebrafish. We tested whether mitochondrially associated Stat3 had a role in PLL axon outgrowth. Cox8 is a subunit of complex IV in the mitochondrial respiratory chain. We co-expressed Stat3 tagged with tdTomato (Stat3-tdTomato) and Cox8a tagged with EGFP (Cox8a-EGFP) in PLL neurons driven by a *5kb-neurod* promoter, and performed sequential imaging of axons at 3 dpf. Fig 4A-B and Video S2 displays a representative result of co-localization of Stat3 and a mitochondrion through time-lapse imaging and kymographs analysis.

**Fig 4.**
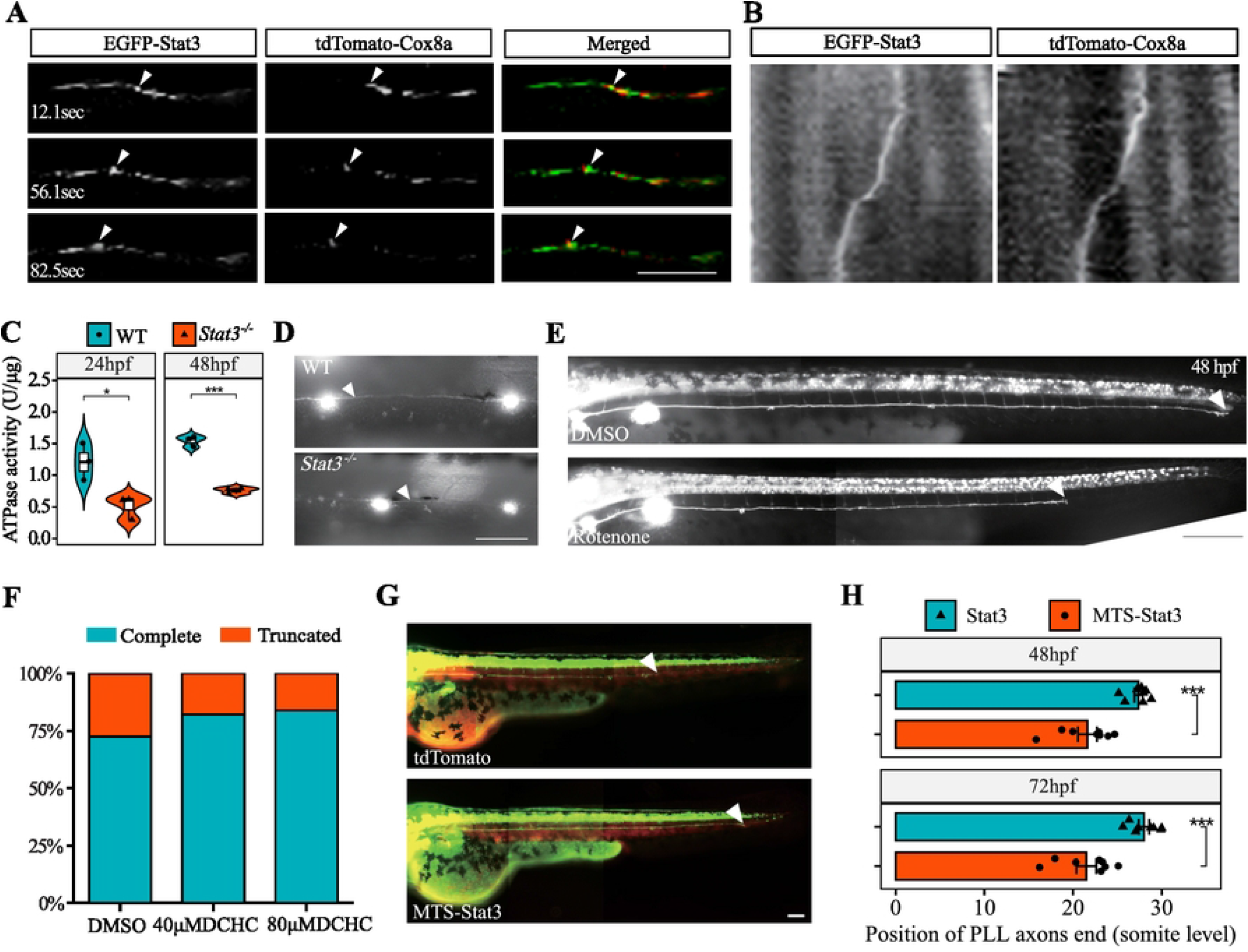
Mitochondrial Stat3 is required for PLL axon growth. (A) A snapshot of video 2 showing the co-transportation of EGFP-Stat3 and tdTomato-Cox8a in PLL axons. Yellow arrows point to a retrograde Stat3/Cox8a positive cargo. (B) Kymographs generated from video 2 for individual cargos. (C) The ATPase activity of wild-type and *stat3* mutant embryo at 24 hpf and 48 hpf. * *p*<0.05 and *** *p*<0.001 (*t*-test), compared with wild-type. (D) Representative TMRE staining images of the PLL neuromasts and axons in wild-type and *stat3* mutant. The arrowheads point to the PLL axons. (E) Representative images of Rotenone- and DMSO-treated embryos under TgBAC*(neurod1:EGFP)* background at 48 hpf. Scale bar: 200 μm. (F) The percentage of truncated versus full length PLL axons of the *stat3* heterozygous mutant incross offspring treated with 40 μM DCHC, 80 μM DCHC or DMSO. (G) The PLL axons of a *stat3* mutant embryo rescued with mitochondria targeting sequence tagged Stat3 (MTS-Stat3) or control (tdTomato) mRNA. Arrow indicates the growth cone of PLL axons. Scale bar: 100 μm. (H) Quantification of PLL axons length of *stat3* mutant rescued with MTS-Stat3 or tdTomato mRNA. *** *p*<0.001 (Wilcoxon RS test) compared with control group.

To investigate the function of *stat3* in zebrafish mitochondria, we measured the ATPase activity of the whole embryos at 24 hpf and 48 hpf, respectively. Notably, the ATPase activity of *stat3* mutants was significantly lower than wild-type at 24 hpf when the PLL axons initiate growth (*N*=3). At 48 hpf, after the PLL axon migration finished, the level of ATPase activity in *stat3* mutant was still reduced (*N*=3) (Fig 4C). Thus, Stat3 is required for mitochondria to maintain ATPase activity during zebrafish embryonic and larval stages. To determine whether the mitochondrial activity in PLL axons was also affected by the *stat3* mutation, the larvae were stained with Tetramethylrhodamine ethyl ester (TMRE) at 48 hpf, which is used to quantify the mitochondrial membrane potential in the PLL ganglion *in vivo* (Mandal et al., 2018). The membrane potential of mitochondria in PLL axons in *stat3* mutant was significantly decreased compared to wild-type (Fig 4D). Thus, Stat3 is involved in the regulation of intermembrane potential of mitochondria in PLL axons.

To confirm the role of mitochondrial activity on PLL axon growth, we treated the zebrafish larvae with 50 μM rotenone, an inhibitor of complex I of the mitochondrial respiratory chain, during PLL axon growth (24 hpf-48 hpf). The PLL axon growth was obviously inhibited by rotenone (Fig 4E). This result was similar to the phenotype in *stat3* mutants. In addition, we treated the progeny of a *stat3* heterozygous mutant incross with DCHC, a promoter of mitochondria activity, and found that exposure to DCHC at 40 μM or 80 μM increased the proportion of complete PLL axons (82.3% and 83.9% vs 72.5%) (Fig 4F; N=120). These results suggest that Stat3 regulation of PLL axon extension is dependent on mitochondria activity.

In mice, Stat3 fused with a mitochondrial targeting sequence (MTS-Stat3) promotes axon regeneration in RGC (Luo et al., 2016). We injected MTS-Stat3 mRNA in *stat3* mutant embryos. MTS-Stat3 mRNA rescued the PLL axon growth in *stat3* mutants with most of the PLL axons reaching to tail, whereas the tdTomato mRNA couldn’t promote PLL axon growth in *stat3* mutants (Fig 4G-H; N=8). Combined with result of NLS-Stat3 mRNA above, we concluded that the mitochondrial Stat3 is sufficient to promote PLL axon growth in zebrafish.

### 5. Jak/Stat signaling is necessary for axon regeneration

We also analyzed PLL axon regeneration in wild-type and *stat3* mutants. The axons were ablated between the fourth and sixth somites at 48 hpf, when the PLL axons completed their embryonic growth. The regenerated axon in both wild-type and *stat3* mutants extended through the ablation site 12 hours post-lesion (hpl). At 48 hpl, PLL axons in wild-type were nearly completely regenerated, but PLL axons in *stat3* mutant failed to regenerate to their original length (Fig 6A). We quantified the length of PLL axons at 20 hpl, 30 hpl, 48 hpl, and 72 hpl and found that PLL axons in *stat3* mutants at the longest timepoint only regenerated to 30.5-39.9% of their original length, and were significantly lower in length than wild-type at every time point (Fig 6B).

**Fig 5.**
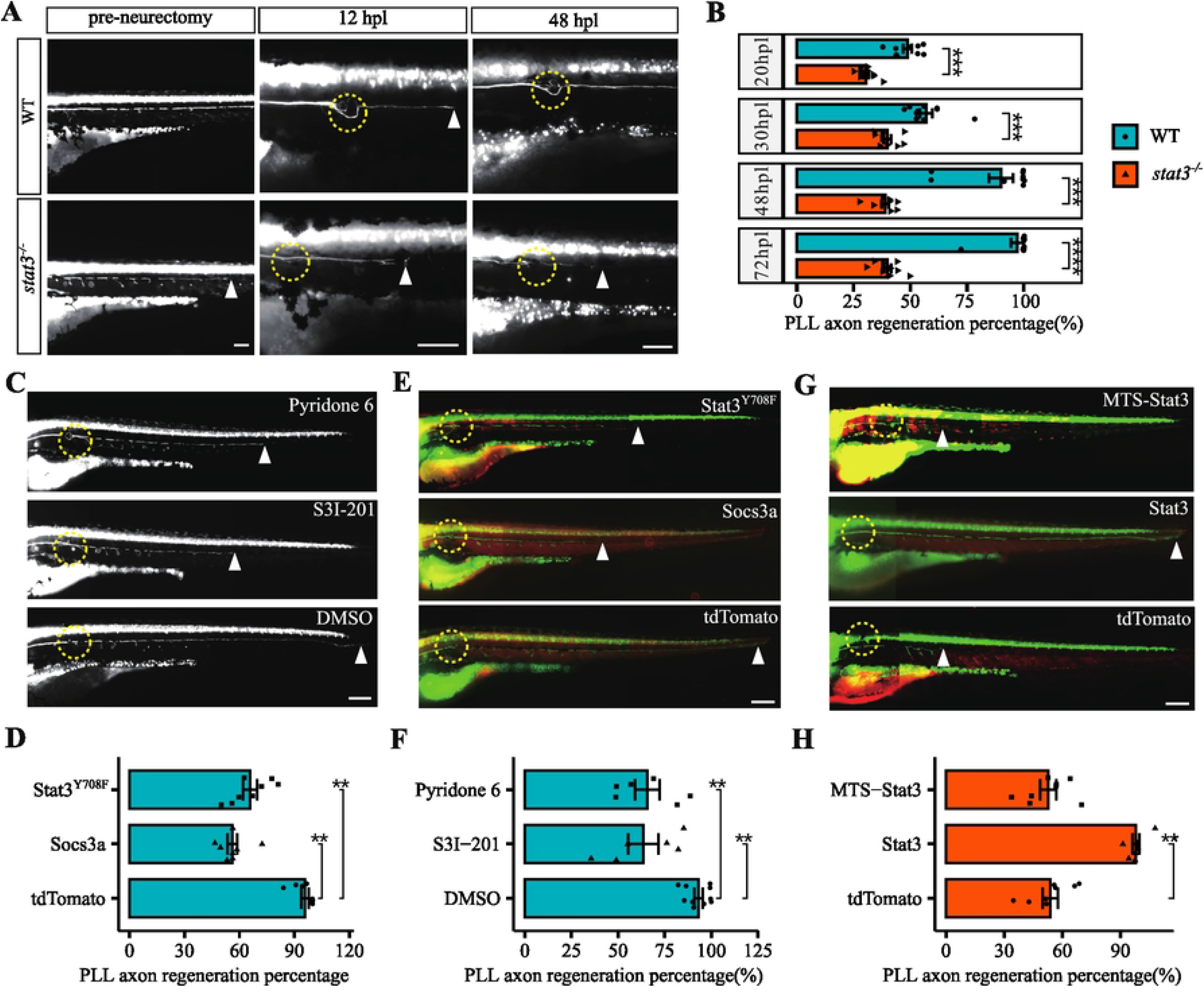
Jak/Stat signaling is essential for PLL axon regeneration. (A) The regeneration of PLL axons in a wild-type and *stat3* mutant at 12 hpl (hour-post lesion) and 48 hpn. (B) Quantification of percentage of PLL axon regeneration in wild-type and *stat3* mutants. (C) The representative image of PLL axon regeneration treated with S3I-201, Pyridone 6, and DMSO (control). (D) Percentage of PLL axon regeneration after Jak/Stat inhibitor treatment. (E) The PLL axon regeneration of wild-types injected with Stat3^Y708F^ (Y708F), Socs3a, or tdTomato (control) mRNA. (F) Axon regeneration percentage shown in (E). (G) The PLL axon regeneration of a *stat3* mutant rescued with MTS-Stat3 and tdTomato (control) mRNA. (H) PLL axon regeneration percentage shown in (G). Yellow circles represent the axon lesion sites. Arrowheads show the terminal of PLL axons. Scale bars: 100 μm. Adjusted *p*-value symbol: \*\**p*<0.01, \*\*\**p*<0.001 and \*\*\*\**p*<0.0001 (Wilcoxon RS test), compared with control group.

**Fig 6.**
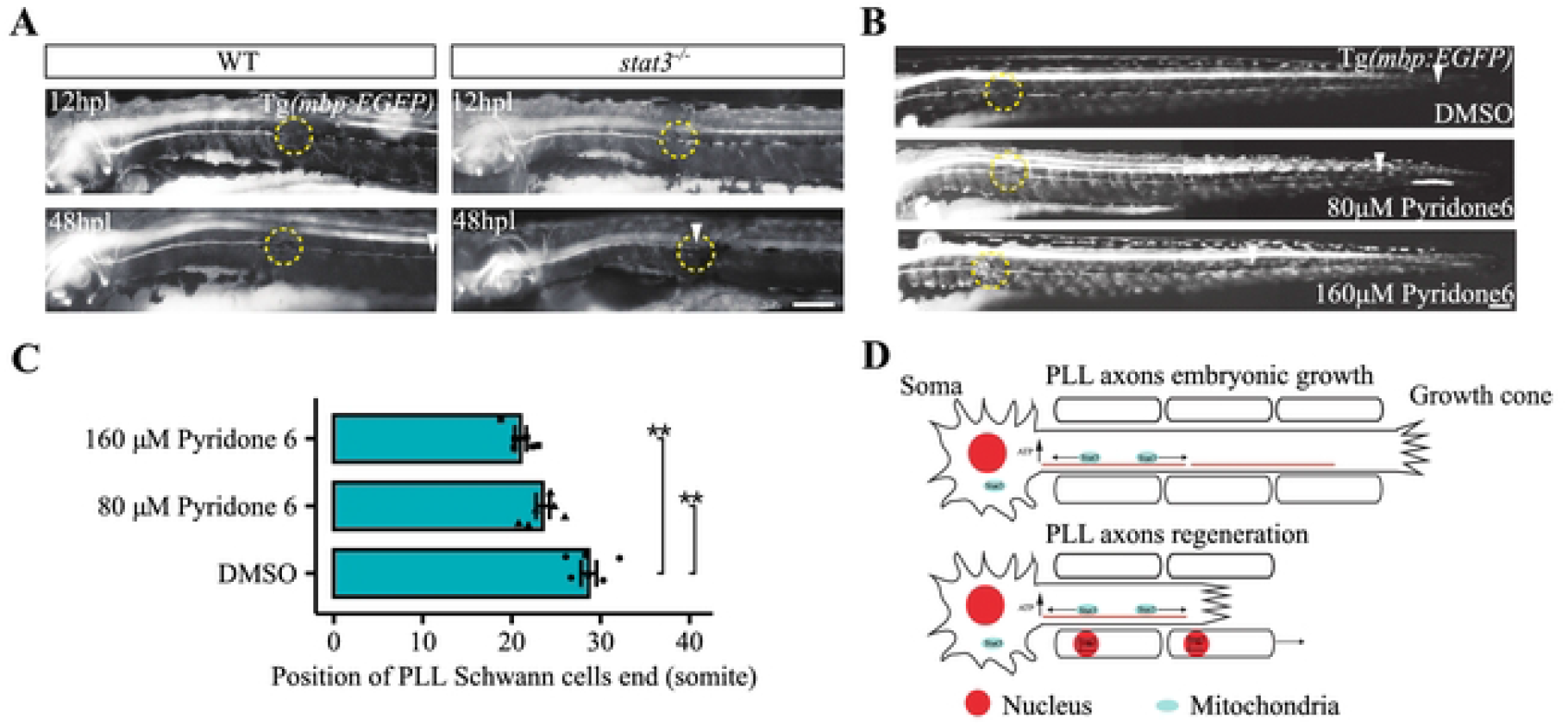
Jak/Stat signaling is involved in Schwann cell migration during PLL axon regeneration. (A) The Schwann cells surrounding PLL axons of wild-type and *stat3* mutants during axon regeneration. Yellow circles represent the axon lesion sites. (B) Representative image of PLL axon regeneration treated with 80 μM or 160 μM Pyridone 6, or DMSO. Arrowheads in (A) and (B) point to the terminus of visible PLL Schwann cells shown by Tg*(mbp:EGFP)*. Scale bars: 100 μm. (C) The quantification of the length of myelination along the length of PLL axons. **adjusted *p*<0.01 (Wilcoxon RS test), compared with control group. (D) Proposed model for how Stat3 regulates PLL axon embryonic growth and regeneration.

To investigate whether Jak/Stat signaling is involved in PLL axon regeneration, we ablated the PLL axons of the wild-type embryos at 48 hpf, then incubated the embryos with Jak/Stat signaling inhibitors. The PLL axons in larvae treated with Pyridone 6 and S3I-201 only regenerated to approximately 65.6% or 63.5% of their original length by 48 hpn, respectively (N=6) (Fig 6C-D). We also injected modified *stat3* mRNA into embryos to generate a dominant-negative effect on Jak/Stat signaling. At 48 hpl, PLL axons in wild-type larvae injected with Stat3^Y708F^ or Socs3a mRNA only regenerated 65.8% or 56.1%, respectively (N=6) (Fig 6E-F). Together, it is suggested that Jak/Stat signaling is required for PLL axon regeneration suggesting there is a separate role for Stat3 in axon regeneration that is not part of normal developmental outgrowth of the PLL axons.

To investigate whether mitochondrial Stat3 was involved in PLL axon regeneration, we injected wild-type Stat3 or MTS-Stat3 mRNA in *stat3* mutants to trigger PLL axons extending to the caudal neuromasts and then ablated the PLL axons at 48 hpf. At 48 hpl, the PLL axons of larvae injected with Stat3 could regenerated nearly completely (97.9%), but the larvae injected with MTS-Stat3 only regenerated approximately 50%, no significant difference from the control (tdTomato) mRNA (N=8) (Fig 6G-H). Taken together, the above results indicate that Jak/Stat signaling is required for PLL axon regeneration, but mitochondrial Stat3 is not sufficient to promote PLL axon regeneration.

### 6. Jak/Stat3 is required for Schwann cell migration during PLL axon regeneration

During regeneration, Schwann cells migrate to the ablation site and guide the regrowth of axons (Min et al., 2021; Villegas et al., 2012). We examined whether Stat3 regulated Schwann cells migration during PLL axon regeneration using the Tg*(mbp:EGFP)* fish line. In wild-type, Schwann cells near distal regions of PLL axons dispersed at approximately 12 hpl, and migrated back to the distal region at about 48 hpl. In *stat3* mutants, the Schwann cells dispersed normally at 12 hpl, but they could not bridge the gap and migrate back to the distal regions of PLL axons (Fig 7A). To determine whether Jak/Stat signaling was involved in Schwann cells migration, we treated the larvae with Jak/Stat signaling inhibitor Pyridone 6. Although the Schwann cells migrated across the ablation gap, Schwann cell migration was significantly inhibited by Pyridone 6 at 48 hpl. The length of Schwann cell surrounding the PLL axons was shorter in larvae treated with 160 μM Pyridone 6 than larvae treated with 80 μM (Fig 7B-C). The inhibition effect of Pyridone 6 on Schwann cell migration was in a dose-dependent manner. Taken together, Jak/Stat3 signaling also plays an important role in Schwann cell-mediated PLL axon regeneration.

## Discussion

In this study, we generated a zebrafish *stat3* mutant line using CRISPR/Cas9 and found *stat3* plays distinct roles in PLL axon growth at the embryonic and regenerative stage. The mitochondrial Stat3 is required for embryonic axon growth through promoting ATP synthesis. However, Stat3 is involved in PLL axon regeneration mainly through promoting Schwann cells migration by Jak/Stat signaling (Fig. 7D). Our findings show a new mechanism of Stat3 in axon growth and regeneration and suggest that Stat3 is an compromising target for axon regeneration.

As a broadly used transcription factor, Stat3 has been found to be essential for embryonic development. Targeted disruption of *stat3* in mice leads to embryonic lethality, which is explained as a consequence of the failure in establishing metabolic exchange between the embryo and maternal blood (Takeda et al., 1997). However, no obvious morphological phenotype was identified in zebrafish *stat3* mutant embryos in previous studies. Maternal and zygotic *stat3* mutants exhibited transient and mild cell cycle increases during zebrafish embryogenesis (Liu et al., 2017). In this study, we found *stat3* was expressed in the PLL ganglion dynamically, and identified a PLL axons truncation phenotype in zebrafish *stat3* mutant embryos. This finding expands our understanding of the function of *stat3* gene in vertebrate embryonic development. Axon extension in Drosophila embryos also requires *stat92E* (Li et al., 2003). It suggests that the Stat family, specifically Stat3, plays critical and conserved role in embryonic axon growth in insects and vertebrates. As reported, blocking *stat3* with antisense morpholinos results in a CaP axon pathfinding defect (Conway, 2006). However, we didn’t detect an obvious phenotype in spinal axons in *stat3* mutant. The phenotype of spinal axons might be an off-target effect of *stat3* morpholino in zebrafish embryonic development. Liu and colleagues found that zebrafish mutants lacking maternal and zygotic Stat3 expression do not exhibit abnormal convergence movements during gastrulation as shown in *stat3* morpholino studies (Liu et al., 2017). Another explanation is genetic compensation from other Stat genes could be occurring in the CRISPR generated allele (Rossi et al., 2015). Stat3 and Stat5 proteins can bind to the same regulatory loci and exhibit functional redundancy to some extent (Wingelhofer et al., 2018). Further studies will be necessary to determine if other members of the Stat family can contribute to embryonic axon extension.

Mitochondria play important roles in axon growth (Sheng, 2017; Smith and Gallo, 2018). As reported, a fraction of Stat3 protein localizes in mitochondria and increases the efficiency oxidative phosphorylation (Wegrzyn et al., 2009). The mitochondrially localized Stat3 has been shown to be important in both cancer cells and heart muscles (Gough et al., 2009; Wegrzyn et al., 2009). Axon growth is an energy consuming process. We showed that Stat3 co-localized with the moving mitochondria in the PLL axons. The ATPase activity and ETC decreased significantly in zebrafish *stat3* mutants during PLL ganglion projecting axons. *stat3* is co-expressed with mtDNA transcription genes in the proliferation tissues of zebrafish and *stat3* knock-out impairs normal mitochondrial transcription and cell proliferation (Peron et al., 2021). Rotenone, a specific inhibitor of mitochondrial complex I, has been used to inhibit neurogenesis in rats (Betarbet et al., 2000). Rotenone treatment mimicked the PLL axon growth defect in *stat3* mutants. As reported, MTS-Stat3 promotes RGC axon regeneration in mice and induces neurite outgrowth in the PC12 cell line (Luo et al., 2016; Zhou and Too, 2011). Mitochondrially-targeted Stat3 could rescue PLL axon growth in zebrafish *stat3* mutant but the nuclear-localized Stat3 could not. We speculate mitochondrial Stat3 plays an important role in energy consuming cells, including differentiating neurons. Mitochondrial Stat3 is also involved in mitochondrial Ca^2+^, reactive oxygen species (ROS) production, and mitochondrial gene expression (Rincon and Pereira, 2018). It will be important to establish if other aspects of mitochondrial function are as important as ATP production for axon outgrowth. Phosphorylation at S727 is critical for Stat3 translocation into the mitochondria (Luo et al., 2016; Zhou and Too, 2011). In this study, we found Stat3^Y708F^ and Stat3^EE435-436AA^ could not rescue PLL axon growth in *stat3* mutants. EE434-435 is required for DNA binding, and Y708 and S727 are required for transcription activation of Stat3 (Horvath et al., 1995; Peron et al., 2021). How this relates to the specific functions in the mitochondria to enable axon outgrowth needs to be explored further.

Although many regeneration processes recapitulate embryonic development, some genes exhibit distinct roles during embryonic development or regeneration. Inflammation in particular, plays a much larger role in tissue regeneration than in embryonic development. Jak/Stat signaling downstream of inflammation signals has been shown to be important for promoting tissue regeneration (Fang et al., 2013). Jak/Stat signaling is essential for optic nerve regeneration in mouse and zebrafish, and spinal nerve regeneration in gecko (Elsaeidi et al., 2014; Smith et al., 2009; Zhang et al., 2020). We show an essential role for Jak/Stat signaling in axon regrowth in zebrafish but not in axon extension during normal development. The regeneration of most peripheral axons is linked to Schwann cell migration which guides the injured axon to target the original location correctly (Ceci et al., 2014; Coleman and Freeman, 2010). We found that *stat3* mutations or Jak/Stat signaling inhibitors hampered PLL axon regeneration and Schwann cell migration. The Schwann cell migration defect is earlier and more severe than the axon regeneration defect. It is tempting to speculate that Jak/Stat signaling promotes PLL axon regeneration through Schwann cell migration, although the contribution of mitochondrial Stat3 in axon regeneration cannot be excluded. In contrast, several lines of evidence indicate that Jak/Stat signaling is dispensable for normal PLL embryonic axon growth. Inhibiting Jak/Stat signaling with small molecules, Socs3a, or a dominant mutant form of Stat3 does not affect PLL axon embryonic growth. Nuclear localized Stat3 (NLS-Stat3) didn’t rescue the axon phenotype of *stat3* mutants. In line with our results, blocking Jak/Stat signaling inhibits DRG (dorsal root ganglion) axon regeneration but not axon embryonic extension (Liu and Snider, 2001). Therefore, Stat3 perhaps plays distinct roles in embryonic and regenerative axon extension.

## Materials and Methods

### 1. Zebrafish strains and husbandry

The AB, TgBAC*(Neurod1:EGFP)* (Obholzer et al., 2008) and Tg*(mbp:EGFP)* (Jung et al., 2010) transgenic lines used in this study were purchased from the China Zebrafish Resource Center. Adult zebrafish were maintained at 28.5°C on a 14 h:10 h light/dark cycle, fed with artemia twice per day. Embryos were collected by natural mating with a male to female ratio of 1:1, raised at 28.5°C in blue water and staged according to hours- or days-post fertilization (hpf or dpf) as described previously (Kimmel et al., 1995). All procedures for zebrafish were approved by the Animal Ethics Committee of Shanghai Ocean University.

### 2. Whole-mount in situ hybridization

The templates for anti-sense RNA probe synthesis were produced by PCR from embryonic cDNA. All primers for probe synthesis are listed in S1 Table. The T7 promoter sequence was added to the 5’-end of anti-sense primers. The RNA probes were synthesized using T7 RNA polymerase with DIG RNA Labeling Mix (Roche). The larvae were fixed in 4% formaldehyde at 4°C overnight, dehydrated with 100% methanol and stored at −20°C. Samples were rehydrated into PBST (1xPBS with 0.1% tween-20) gradually, bleached with 10% H_2_O_2_, permeabilized with 10 μg/μL proteinase K for 10 min, then post-fixed in 4% PFA for 20 min. Hybridization was carried out at 70°C overnight, followed by post-hybridization washing, blocking with 2% bovine serum albumin (BSA) and 2% sheep serum, incubation with anti-DIG antibody (Roche), and stained with NBT/BCIP solution (Roche).

### 3. Generation of zebrafish stat3 mutant line

Zebrafish *stat3* mutants were generated by Clustered Regularly Interspaced Short Palindromic repeats (CRISPR)/ CRISPR-Associated Protein 9 (Cas9) according to the published protocol (Varshney et al., 2016). In brief, a single guide RNA (sgRNA) targeting to *stat3* exon 3 (5’-GGTGCTGCTTGATGCGCCGCAGG-3’) was designed using the Zebrafish Genomics track in UCSC Genome Browser (LaFave et al., 2014). The target-specific oligo (S1 Table) including a T7 promoter and the universal oligo with the crRNA-tracRNA sequence were used for producing the template for sgRNA synthesis. The sgRNA and capped Cas9 mRNA were synthesized by *in vitro* transcription. Approximately 1.4 nL of sgRNA (about 50 pg) and capped Cas9 mRNA (about 300 pg) was co-injected into wild-type embryos at the one-cell stage. The genomic DNA was extracted from each injected embryo using alkaline lysis. The insertion and deletion (In-Del) efficiency was evaluated by fluorescent PCR using the *stat3* genotyping primer set (S1 Table), followed with capillary electrophoresis (Carrington et al., 2015). The embryos with high In-Del efficiency were established as founder fish and raised to adulthood. The F1 were obtained by crossing the founder with AB, fin-clipped and genotyped as described above. A pairwise heterozygous F1with selected mutation was incrossed to get the *stat3* mutant.

### 4. qPCR

The embryos derived from a *stat3* heterozygous mutant incross were genotyped by PCR analysis of genomic DNA extracted from embryonic fin clips at 48 hpf and separated according their genotypes. Total RNAs of 20 wild-type or *stat3* mutants was extracted using TRIzol reagent (Invitrogen). cDNA of wild-type and *stat3* mutants were generated by the ProtoScript First Strand cDNA Synthesis Kit (NEB) using random primers. qPCR was performed with the Luna Universal qPCR Master Mix (NEB) on an ABI PRISM 7000 Real-Time PCR System (Applied Biosystems, USA). The *elf1α* gene was used as endogenous control. The qPCR primers for *elf1α, atf3, c-jun, klf6a, klf7a, socs3a, socs3b, tub1a, tub1b, tub1c*, and *thymosin* gene refer to previous study (McCurley and Callard, 2008; Elsaeidi et al., 2014). The qPCR primers for *stat3* and *il6* are listed in S1 Table. All reactions were performed in technical triplicates. Relative expression level of each gene was calculated using 2^-ΔΔCt^ method and normalized to the mean of control group. The results represent biological replicates, including the standard error of the mean.

### 5. Immunofluorescence and TMRE staining

The larvae were fixed in freshly-made formaldehyde as described above. After three washes with PBST for 20 min each, samples were incubated in blocking solution (2% BSA and 2% normal goat serum in PBST) for 1 h, and in mouse acetylated-tubulin antibody (1:1000 dilution, Sigma-Aldrich) at 4°C overnight. After three washes with PBST for 20 min each, samples were incubated in goat anti-mouse IgG conjugated with Alexa Fluor 488 (1:200, Invitrogen) for 2 h at room temperature, then three washes with PBST. Samples were visualized using a Zeiss Axio Observer fluorescent microscope (Carl Zeiss), the genomic DNA of each fixed sample was extracted with the FastPure FFPE DNA Isolation Kit (Vazyme) and the genotype was determined as above.

Zebrafish larvae at 3 dpf were incubated in 2.5 μM mitochondria specific dye tetramethylrhodamine ethyl ester (TMRE, diluted with Holtfreter’s buffer) for 1 h in the dark. After washing with Holtfreter’s buffer twice, the larvae were immobilized in 2% low-melting-point agarose (LMP, Fisher Scientific) and visualized using the 568 nm excitation channels.

### 6. Generation the constructs

All constructs used in this study were generated by seamless cloning strategy using the ClonExpress II One Step Cloning Kit (Vazyme). The Tol2 vector backbone were amplified from pHuC:farnesylated-TdTomato (Faucherre et al., 2009), the full length of the coding sequence (CDS) of *stat3, socs3a*, and *cox8a* was amplified from zebrafish embryonic cDNA, and 5 kb-*neurod* promoter in zebrafish was amplified from zebrafish genomic DNA as described (Drerup et al., 2016). Firstly, the pHuC:TdTomato-stat3 was generated by fusing *stat3* CDS downstream of tdTomato in pTol2-HuC:farnesylated-tdTomato. pNeurod:tdTomato-Stat3 was constructed by replacing the HuC promoter in pHuC:tdTomato-Stat3 with zebrafish 5 kb-*neurod* promoter. pNeurod:tdTomato-Cox8a were generated by replacing *stat3* CDS in pNeurod:tdTomato-Stat3 with zebrafish Cox8a CDS. These constructs were used to express *stat3* or display the mitochondria in PLL axon.

The plasmids for mRNA synthesis were modified from pcDNA3.1. tdTomato was cloned into the multiple cloning site of pcDNA3.1 as a control. The CDS of *socs3a* and *stat3* was fused downstream of tdTomato, respectively. Constructs for expressing mutated Stat3^Y708F^ and Stat3^EE435-436AA^ were generated by modifying pcDNA3.1-Stat3 using Mut Express II Fast Mutagenesis Kit V2 (Vazyme). Similarly, constructs for expressing Stat3 tagged with nuclear localization sequence (NLS) and mitochondrial targeting sequence (MTS) were generated by amplifying the whole pcDNA3.1-Stat3 with primers containing NLS or MTS coding sequence at their 5’-end and re-ligating. All the constructs were verified by sequencing.

### 7. Transmission electron microscopy (TEM)

The wild-type and *stat3* mutant zebrafish was euthanized with tricaine methanesulfonate (MS222, Sigma) and fixed using 2.5% glutaraldehyde (diluted in PBS) at 4°C overnight. After rinsing in 0.1 M sodium cacodylate buffer, the samples were post-fixed in 1% osmium tetroxide for one hour, and then rinsed in 0.1 M sodium cacodylate buffer again. Larvae were then dehydrated with a graded series of ethanol, infiltrated with and embedded in epoxy resin, and polymerized in a 70°C oven for 48 h. After identifying areas of interest, ultra-thin sections of 100 nm thickness for TEM analysis were then cut using an ultramicrotome and stained with 33% methanolic uranyl acetate for 15 minutes and lead citrate (Electron Microscopy Sciences) for 7 minutes. Digital electron micrographs were collected using a HITACHI H-7700.

### 8. mRNA and plasmid injection and quantification of axon growth

The capped mRNA of Tol2 transposase, TdTomato, Socs3a, wild-type, mutant, and tagged Stat3 were synthesized with linearized plasmids using T7 mMessage mMachine kit (Invitrogen). For overexpression essays, one nanogram of the capped mRNA was injected into the cytoplasm of each embryo derived from *stat3* heterozygous mutant incross, with *TgBAC(Neurod1:EGFP)* background, respectively. To get the transient transgene, the mixture of Tol2 plasmid with Tol2 transposase mRNA was used for injection. Following injections, the PLL nerves of each larva were visualized and quantified according the position of PLL axon terminal at 2 dpf. And each larvae injected was genotyped as above.

### 9. Drug treatments

In this study, inhibitors of Jak/Stat signaling, mitochondrial respiratory chain, and mitochondrial boosters were used to treat zebrafish embryos. All compounds were dissolved in DMSO and stored at −80°C in small aliquots. Respectively, the embryos of Tg*BAC(Neurod1:EGFP)* were treated with 80 μM Pyridone 6 (MCE, HY-14435), 400 μM S3I-201 (MCE, HY-15146), 5 μM Stattic (MCE, HY-13818), and 50 nM rotenone (MCE, HY-B1756) diluted with Holtfreter’s buffer from 24 to 48 hpf. Equal volume of DMSO was used as control. The length of PLL axons of the treated larvae was quantified based on the somite position. The progeny of *stat3* heterozygous mutant incrosses using the TgBAC*(neurod1:EGFP)* background were treated with 50 μM DCHC (Sigma, D5569) from 24 to 48 hpf. The percentage of PLL axons reaching the tip of tail was calculated. PLL axons of TgBAC*(neurod1:EGFP)* or Tg*(mbp:EGFP)* background were amputated between the 3^rd^ and 8^th^ somites using electric neurectomy methods at 3 dpf as described previously (Ceci et al., 2014). Then, the larvae were treated with the Jak/Stat inhibitor, Pyridone 6 (80 μM) for 2 days during PLL axon regeneration. The PLL nerves were imaged using the 488 nm excitation channels and the length of PLL axons were quantified as above.

### 10. ATP synthase activity

ATP synthase activity of wild-type and *stat3* mutant zebrafish embryos was measured using the ATP Synthase Specific Activity Microplate Assay Kit (Abcam) following the manufacturer’s protocol. Briefly, 25 embryos of wild-type and *stat3* mutants were homogenized in 200 μL of homogenization buffer. After freezing and thawing, the homogenates were centrifuged at 16,000 rpm and the solubilized fraction was collected. After adjusting the protein concentration to 5.5 μg/μL, the sample was extracted in Detergent Buffer on ice for 30 min followed by centrifugation at 16,000 rpm for 20 min. Fifty μL of supernatant was loaded in each well of a pre-coated microplate and kept at 4°C overnight. Solution 1 was used as blank reference. Each well was rinsed twice with Solution 1 and incubated with Lipid Mix for 45 min. Reagent Mix was added to the reaction and ATP synthase activity was measured at OD340 at 1 min intervals for 1-2 h. For quantifying the mount of ATP synthase, each well was loaded with Solution A and incubated at room temperature for 1 h. After washing with Solution 1 twice, the wells were loaded with Solution B and incubated at room temperature for 1 h. Development Solution was added to each well, and the amount of ATP synthase was measured at OD405 at 1 min intervals for 30 min.

### 11. Axonal mitochondria localization analysis of Stat3

To display the transport of Stat3 and mitochondria in PLL axons, Neurod:TDT-Stat3 and Neurod:EGFP-Cox8 plasmids together with Tol2 transposase mRNA, were co-injected in the one-cell stage embryos of AB. The embryos expressing EGFP and tdTomato in a single PLL ganglion cell were mounted in 1.5% low melting point agarose, anesthetized with 0.02% tricaine on a glass bottom plate and imaged with 488 nm and 568 nm excitation channels in sequence at 72 hpf. Photographs were collected at fastest speed for 1 min. The mitochondrial transport in PLL axons was analyzed using kymograph analysis in the ImageJ FIJI. Kymographs were generated from each imaging session.

### 12. PLL nerve amputation

Wild-type or *stat3* mutant larvae with the TgBAC*(Neurod:EGFP)* and Tg*(mbp:EGFP)* transgenes were anesthetized in MS-222, and the PLL axons were amputated between the 3^rd^ and 8^th^ somites with an eye scalpel under a dissecting microscope. The larvae after axonectomy were recovered in Holtfreter’s buffer and imaged under fluorescent inverted microscope.

### 13. Statistical analysis

Statistical analysis and graphs were performed using R with the packages “rstatix” and “ggpubr”. Normality of the data was tested using Shapiro-Wilk test. Two groups comparisons were analyzed using Student’ *t* test or Wilcoxon Rank Sum Test as described in the figure legends. For multiple groups comparisons, differences between treatment groups and their matched controls were tested, and adjusted *p*-value were calculated using the False Discovery Rate (FDR, Benjamini & Hochberg) controlling method. Differences of proportions for categorical variables were tested using Fisher’s exact test. For all comparisons, *p* or adjusted *p* <0.05 was considered as statistically significant.

## Acknowledgements

We thank Donghua Zhou, Yuanhang Chen, and Xujing Zhu, who participated in this study for their technical assistance and helpful discussions. We also thank comments and suggestions from Xiaojie Wang. This study was supported by the National Natural Science Foundation of China (NSFC) (Grant 31772406 to C.F. and Grant 31702329 to J.W.).

## Supporting information captions

**S1 Table. Nucleotide sequences of primers used in this study.**

**S1 Fig. The PLL axons did not degenerate in *stat3* mutants.**

(A-B) The PLL axons in wild-type (A) and *stat3* mutants (B) shown using TgBAC*(neurod1:EGFP)* transgenic background at 17 dpf. The scale bar is 25 μm.

**S2 Fig. The number of cell body in PLL ganglion was not changed significantly in *stat3* mutants.**

(A) The PLL ganglion in wild-type and *stat3* mutants shown using TgBAC*(neurod1:EGFP)* transgenic background at 3 dpf. The scale bar is 100 μm. (B) Quantification of the PLL ganglion cell body in wild-type and *stat3* mutants.

**S3 Fig. The DLL axons in *stat3* mutants was shorter than wild-type.**

(A) The DLL ganglion in wild-type and *stat3* mutants shown using TgBAC*(neurod1:EGFP)* transgenic background at 3 dpf. Arrow head indicates the tip of DLL axons. (B) Quantification of length of DLL axons in wild-type and *stat3* mutants. (C) The spinal cord axons in wild-type and *stat3* mutants shown using TgBAC*(neurod1:EGFP)* transgenic background at 5 dpf. Arrow head indicates the tip of spinal cord axons. Arrow indicates the branch of spinal cord axons. The scale bars in (A) and (C) are 100 μm.

**S4 Fig. qPCR analysis of the effect of stat3 mutation on expression of genes associated with regeneration.**

\**p*<0.05, \*\**p*<0.01, \*\*\**p*<0.001 and \*\*\*\**p*<0.0001 (*t*-test), compared with wild-type.

**S1 Video. The time-lapse of PLL axons growth in a wild-type and *stat3*^-/-^ from 24 hpf to 40 hpf.**

The embryo is oriented with head to the left, dorsal up. Time is in minutes.

**S2 Video. Stat3 and Cox8a labeled mitochondria transport in the PLL axons.**

The embryo is oriented with head to the left, dorsal up. Time is in seconds.

## Notes

### Competing Interest Statement

The authors have declared no competing interest.

